# Scale-resolved analysis of brain functional connectivity networks with spectral entropy

**DOI:** 10.1101/813162

**Authors:** Carlo Nicolini, Giulia Forcellini, Ludovico Minati, Angelo Bifone

## Abstract

Functional connectivity is derived from inter-regional correlations in spontaneous fluctuations of brain activity, and can be represented in terms of complete graphs with continuous (real-valued) edges. The structure of functional connectivity networks is strongly affected by signal processing procedures to remove the effects of motion, physiological noise and other sources of experimental error. However, in the absence of an established ground truth, it is difficult to determine the optimal procedure, and no consensus has been reached on the most effective approach to remove nuisance signals without unduly affecting the network intrinsic structural features. Here, we use a novel information-theoretic approach, based on von Neumann entropy, which provides a measure of information encoded in the networks at different scales. We also define a measure of distance between networks, based on information divergence, and optimal null models appropriate for the description of functional connectivity networks, to test for the presence of nontrivial structural patterns that are not the result of simple local constraints. This formalism enables a scale-resolved analysis of the distance between an empirical functional connectivity network and its maximally random counterpart, thus providing a means to assess the effects of noise and image processing on network structure.

We apply this novel approach to address a few open questions in the analysis of brain functional connectivity networks. Specifically, we demonstrate a strongly beneficial effect of network sparsification by removal of the weakest links, and the existence of an optimal threshold that maximizes the ability to extract information on large-scale network structures. Additionally, we investigate the effects of different degrees of motion at different scales, and compare the most popular processing pipelines designed to mitigate its deleterious effect on functional connectivity networks.

## 1. INTRODUCTION

Complex networks theory provides a robust framework to study the structural and functional organization of brain connectivity, which can be naturally represented as a graph, a collection of nodes (anatomical brain regions) and edges (functional or structural coupling between nodes) [1] (Figure 1).

**Figure 1.**
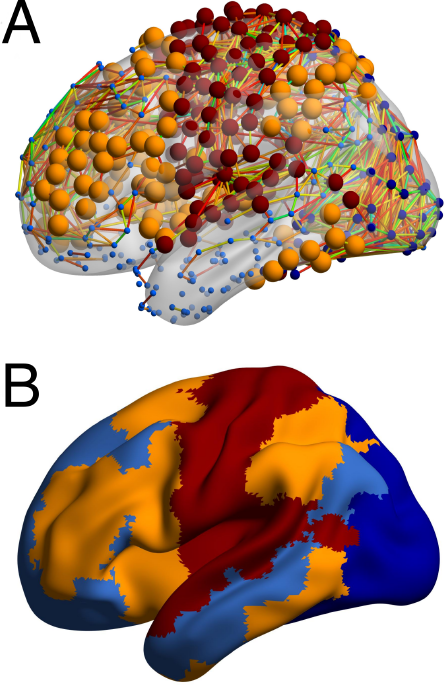
Examples of a resting state functional brain network. In Panel A., nodes represent brain regions, connected by links whose value is obtained by averaging Fisher transformed Pearson correlations over many subjects. Radius of nodes indicates the degree, while their colors indicate the modular membership as overlaid on a surface template in Panel B. The partition entails four communities and is found with the Louvain algorithm [13], modularity value *Q* ≈ 0.4.

Several foundational concepts have been borrowed from network theory and have become part of the neuroscience parlance. By way of example, properties like small-worldness [2], scale-freeness [3] and modularity [4–6] have been demonstrated in brain networks, providing insight into their complex topological organization and its bearing on the dynamical processes underlying brain function in health and disease [7].

However, despite its increasing popularity, this approach is still the subject of debate, and a number of seemingly simple, yet critically important questions remain open. For example, a widely accepted measure of the distance between graphs, e.g. to compare empirical networks with an appropriate null model, is still missing, and so is a measure of the information encoded in a network. This problem is exacerbated by the lack of a ground truth structure for empirical brain networks.

Neuroimaging methods, like functional MRI (fMRI), electroencephalography and magnetoencephalography (EEG, MEG) are often used to assess brain functional connectivity; the resulting networks depend tremendously on data processing and experimental parameters, but the lack of an established reference makes it challenging to determine the optimal procedures univocally. Glaring examples of this problem include the ongoing debate regarding the use of Global Signal Regression [8] in resting-state functional connectivity, or the application of thresholds for network sparsification [9–11].

Specific network metrics (such as node degree distribution or modularity) have often been used for network comparison, but they do not capture the intrinsically multiscale structure of brain networks, focusing only on specific local or global features. Moreover, such measures describe properties that may also be present in random networks. By way of example, large values of modularity have been observed in random networks [12] as well as in natural networks. Hence, a measure of the distance between a given brain network and an appropriate null model, i.e., a random counterpart that satisfies certain constraints, would be essential to address the fundamental question: how far from random are brain networks?

Information-theoretic measures based on the Shannon entropy would appear to be a natural choice to address this issue, but an extension of this formalism to complex networks has proven challenging, although useful in specific contexts [14, 15].

Recently, pioneering work by De Domenico and Biamonte [16] has demonstrated the use of spectral entropies to define distances between pairs of complex networks. Specifically, those authors recognized that the Laplacian of the adjacency matrix describing a given network can be used to construct a matrix that satisfies the same mathematical properties of quantum mechanical density matrices, thus enabling the extension of von Neumann entropy to complex networks. Several implications of this elegant development may be useful for the analysis of brain networks. Differently from other methods, this approach does not rely on a subset of network descriptors [17], but provides an information-based measure that takes into account the entire network structure at all scales. The strength of this formalism lays in the dynamical description of a diffusion process taking place over the network, with the characteristic time described by a parameter *β* that determines the diffusion scale. Hence, spectral entropies provide a scale-resolved, information-based metric to define and optimize network models.

The same framework enables measuring the distance between networks that can be rigorously defined in terms of quantum relative entropy, or information divergence [18]. This quantifies the information gain when a model is used to explain an empirical observation. Moreover, minimization of relative entropy can be used to optimize model parameters [16, 19], or to select different models based on their ability to reproduce the data.

Here, we extend the novel formalism to the study of brain functional connectivity, and demonstrate its potential to address a few open problems in the analysis of resting-state connectivity networks.

Firstly, we implement two new models of maximally-random networks with specific local or global properties to evaluate their deviation from empirical counterparts at different scales. We also propose computationally efficient procedures to fit them to empirical networks. The simpler model describes a class of networks where the total number of links and total weight are constrained to match those of the empirical network. The latter model addresses the more general case in which both degree and strength sequence are preserved upon randomization of the edges. As an important point of novelty, these models are applicable to networks with continuous weights, like functional connectivity networks, and can be optimized constraining their density to that of the empirical network.

A most contentious methodological issue in graph analysis, as applied to the study of brain connectivity, is the one of network sparsification. Functional connectivity networks are generally derived from pairwise correlations of spontaneous fluctuations extracted from each pair of brain regions, resulting, by definition, in a fully connected weighted matrix. However, dense networks are computationally demanding, and weak links, which represent the overwhelming majority of edges, might contain spurious correlations. Network sparsification is then an essential step to recover the network structure.

A number of different thresholding techniques have been proposed [9–11, 20–22] but the choice of the threshold, which strongly affects the topological features extracted from the network, remains somewhat arbitrary. Adding to this, it is now argued that the sparsification procedure itself might actually inject artifactual structures within the network of interest. A recent study, indeed, revealed the introduction of some complex features in random networks as a sole result of thresholding [23]. Altogether, given the lack of agreement on the best thresholding level, and the uncertain significance of the weakest and negative links, there is a trend to completely avoid the application of a threshold, and to work with fully connected networks [24–27]. Starting from the above considerations, here we apply the novel formalism of spectral entropies to investigate the effects of thresholding on resting-state networks, with the aim to assess whether this practice is beneficial for the extraction of information on the large-scale organization of the network, and whether an optimal threshold that maximizes separation between the empirical network and its random counterpart exists.

Other open issues in the functional connectivity field relate to more basic aspects of image processing. For example, it is well known that small head movements during the resting-state fMRI scan can substantially impact the subsequent functional connectivity analysis [28–30]. Motion artifacts are a significant cause of spurious correlations that can substantially affect the structure of functional connectivity networks derived from functional MR. The search for an optimal strategy for the correction of motion-related noise has become a center of attention in the field [31, 32]. Moreover, non-neural physiological activity, like cardiovascular pulsation, respiratory cycle and autonomic fluctuations can inject spurious correlations across multiple frequencies [33, 34]. A plethora of different noise correction techniques have been introduced, all aiming at the reduction or removal of the impact of in-scanner motion effects [31, 35]. However, a consensus on the most effective approach to removing motion effects without substantially affecting the network intrinsic structural features is still lacking. For example, a major debate revolves around the application of a global signal regression (GSR) [36, 37]. The aggressiveness of approaches based on GSR, and the subsequent introduction of negative correlations in the network following its application, has made it the object of concern in the neuroscientific community, despite its apparent efficacy in removing spatially correlated spurious fluctuations. Here, we seek to demonstrate the potential use of the spectral entropy formalism to study the effects of some of the most popular motion correction procedures on network structure. To this end, we assess the effects of motion on the information contained in the network at different scales, and compare the efficacy of various data processing pipelines for the recovery of structural information in the presence of different degrees of motion.

## 2. MATERIALS AND METHODS

### 2.1. Theoretical framework

In this theoretical section we introduce the notation and describe the formalism of maximum entropy random graph models that is central to this manuscript. Firstly, we describe it in the context of classical entropy, as a means to fit network models to empirical networks. We then introduce two null models, the *Continuous Weighted Thresholded Enhanced Random Graph Model* (CWTERG), a real-valued version of the Erdős-Renyi model, and the *Continuous Weighted Thresholded Enhanced Configuration Model* (CWTECM), which also includes constraints on the node degree distribution. Finally, we introduce the formalism of spectral entropy and define a rigorous measure of network distance based on spectral relative entropy.

We summarize here a few definitions that are necessary to make this paper self-contained. We consider undirected weighted graphs *G* = (*V*, *E*) with |*V*| = *N* number of nodes, |*E*| = *L* number of links and *W* total edge weight. We denote the weighted adjacency matrix as **W** = {*w*_*ij*_}, the binary adjacency matrix **A** = *a*_*ij*_ and the weighted graph Laplacian as **L** = **D** − **W**, where **D** is a diagonal matrix of the node strengths. We indicate the degree and strength as *k*_*i*_ = ∑_*i≠j*_ *a*_*ij*_ and *s*_*i*_ = ∑_*i≠j*_ *w*_*ij*_, respectively. The Heaviside step function is indicated as Θ(*x*).

### 2.2. Classical Maximum entropy random graph models

We let **G** denote a network in a random graph ensemble 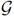, and **G\*** an observed empirical network. The ensemble 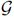 consists of all networks with the same number of nodes *N* and of the same type (undirected, weighted etc.) as **G**, including **G\*** itself. Our goal is to find an analytical description of the random graphs **G\*** that share the same network descriptors of **G\***, and to eventually be able to sample networks from the ensemble. In other words, we look for the functional form of the probability distribution *P*(**G**) over the ensemble 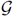, for which the values of descriptors are on average as close as possible to those of the empirical network. We denote the chosen descriptors by **C\*** = **C**(**G\***). These are network-related quantities, like the number of links, the total weight, or the node and strength sequences, and are instrumental in shaping the analytic form of the ensemble. By standard probability arguments, the expected value of the descriptors **C**(**G**) over the ensemble 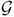 are found as

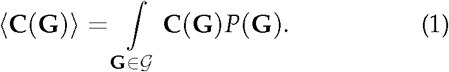

The functional form of *P*(**G**) can be obtained by Shannon entropy maximization subjected to the constraints represented by **C**. This procedure is rooted in Jaynes’s Maximum Entropy formalism [38], a statistical mechanics principle that leads to exact expressions for the probability of occurrence of any graph model. A standard derivation [17, 39] shows that the solution of constrained entropy maximization problem is found by introducing a vector of Lagrange multipliers ***θ***, one for each of the constraints in **C**. The resulting conditional probability reads:

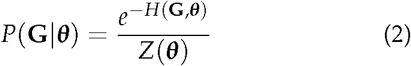

where *H*(*G*, ***θ***) is the graph Hamiltonian, defined as a linear combination of constraints:

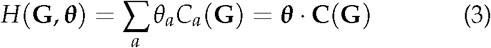

and the denominator *Z*(***θ***) is a normalizing quantity called *partition function*, defined by marginalization over all networks **G** in the ensemble 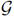:

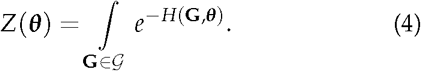

The above results show that the graph probability *P*(*G*|***θ***) depends on the Lagrange multipliers ***θ***, and that it is a function of the constraints considered.

For model fitting purpose, it can be shown [17] that the log-likelihood

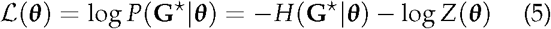

is maximized by the particular value ***θ***\* such that the ensemble average ⟨*C*⟩_***θ***_* of each constraint equals the empirical value *C*(**G**\*) measured on the real network:

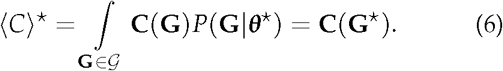

For maximum-entropy ensembles, the maximum likelihood principle indicates the choice of parameters that meet the constraints, and defines a procedure for model fitting: either by maximizing the log-likelihood from Eq. 5 by means of gradient based numerical optimization methods [40], or alternatively by solving the system of nonlinear equations defined by Eq. 6.

In the following, we show a practical application of this approach to a class of null models suitable for the description of resting-state brain connectivity.

### 2.3. Null models for continuous (real-valued) thresholded networks

In network science, and in the study of statistical properties of graphs, a null-model is a mathematical entity representing a family of graphs that match some of the properties of a network, while remaining maximally non-committal with regard to properties not explicitly specified [17, 38, 39]. Null models provide a powerful way to test whether observed nontrivial structural patterns are emerging from simpler local constraints, or they are genuinely present in the empirical network.

Desirable null models do not trade complexity for sufficiency or redundancy [41]: two constraints leading to the same ensemble should be merged into one. Moreover, null models should be neither too complicated nor too simple. Too many parameters and the null model no longer represents the state of maximal agnosticism as overfitting the data precisely displays all its features. Conversely, too few parameters and the picture it conveys is oversimplified, hence lacking explanatory power.

In this sense, we are looking for a model of rs-fMRI networks that is complex enough to match simple local features of the network, but remaining completely uninformative over higher-order patterns. Local constraints such as the number of links or the degree sequence may already fully contain all the information conveyed by the network. Networks of this kind have no statistical patterns or regularities [41, 42] beyond those described by local properties: local features explicitly enforced represent a null hypothesis that we can use as a reference to quantify significant deviations or patterns.

In designing a null model for resting-state fMRI networks, we should take into account the continuous nature of link weights, the binary backbone and weighted structural patterns.Importantly, here we deal only with positive link weights.

Here we extend previous random graph models [43–46] to networks with real-valued links distributed over a connectivity backbone modeled by the degree and strength sequence. These local variables are the optimal trade-off to shape the irreducible and unavoidable complexity needed to accommodate the heterogeneous structure of real networks. Nonetheless, we also describe a simpler model with only two global constraints, namely the total number of links and weight. In the next section we will show that the the former model captures most of the long-range connectivity and mesoscopic structure, while the latter describes adequately local features but remains uninformative of larger-scale structure. Hereby, we embrace the Exponential Random Graph formalism [1, 39] and analytically build the maximally random counterpart of empirical networks where only specific features are reproduced, on average. We first describe the model where the number of links and total weight are constrained and then move to the more general case where both the degree and the strength sequences of the network are considered. Most importantly, both models include a hard thresholding procedure [10, 22, 47] that is often used in analysis pipelines.

#### 2.3.1. Random networks with fixed links number and weight

We introduce the random graph model that fixes the average total number of links *L\** and the average total weight *W\**, together with an external threshold parameter *t* with the name of *Continuous Weighted Thresholded Enhanced Random Graph Model*. This model is obtained by a Hamiltonian that explicitly enforces these two constraints:

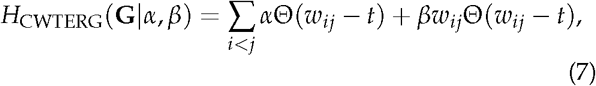

where the Lagrangian multipliers ***θ*** of the problem are the two scalars, *α* and *β*. This Hamiltonian is designed to weight the contribution of binary links with the term *α* and the contribution of weighted links with the term *β*. The role of the threshold parameter *t* becomes clear if a dense network is fed in the model, and its null network is sought for as a function of the threshold. Degrees of a network are sum of binary variables, and the Heaviside function Θ is exactly centered at *t*, taking values one or zero if the edge weight exceeds the cut-off threshold. Similarly, the threshold *t* shapes the sequence of nodes strength, by contributing with a factor ∑_*j*_ *w*_*ij*_ for weights greater than the cut-off *t*. For notation clarity, a change of variables can be performed and the original Lagrangian multipliers are replaced by their exponentiated counterparts, namely the variables *x* = *e*^−*α*^ and *y* = *e*^−*β*^.

The partition function *Z*_CWTERG_ is obtained from the marginalization over all networks in the ensemble as in Eq. 4. A simple calculation for this case (see ref. [17]) yields:

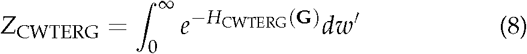

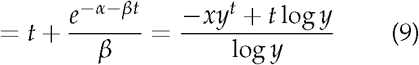

The expected number of binary links is found by taking the derivatives of the free energy [39], *F* = − log*Z* with respect to *α*, the Lagrangian multiplier pertaining the binary links. Similarly, the expected total weight is the derivative with respect to *β* of the free energy. As a result for the CWTERG we get the expressions for the link probability and expected weight, relatively:

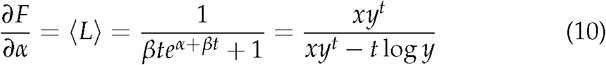

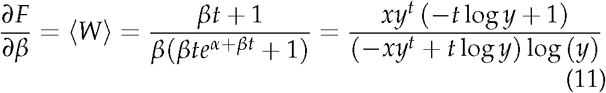

Fitting the CWTERG model to empirical networks requires one to simultaneously solve a system of two nonlinear equations, and finding the values of the Lagrangian multipliers *x*, *y* such that:

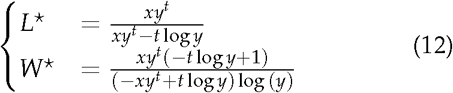

Alternatively, and in a completely complementary fashion, one can maximize the log-likelihood of the model 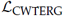, calculated as the logarithm of the conditional probability *P*(**G**|*x*, *y*):

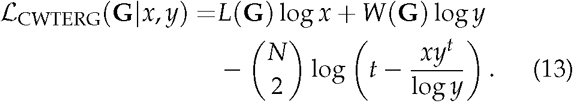

#### 2.3.2. Random networks with fixed degrees and strengths

The CWTERG model describes the ensemble of networks whose total weight and number of links are constrained to some empirical values. Hence it can be considered an extension of the Erdős-Renyi random graph model to thresholded weighted networks. However, this model only describes networks with uniform connectivity patterns, as it is not considering the heterogeneity of the degrees and strengths.

The Continuous Weighted Thresholded Enhanced Configuration Model (CTWECM) overcomes this problem by defining a Hamiltonian

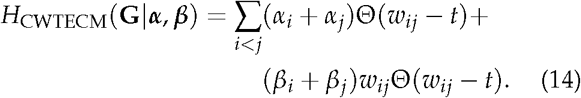

where *α*_*i*_, *β*_*i*_ are the Lagrangian multipliers. The structural form of the Hamiltonian of the CWTECM is the same as the one of the CWTERG, but now the probability *P*(**G**|***α***, ***β***) can be factorized over all pairs of nodes as follows:

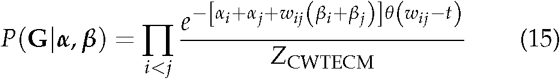

where here 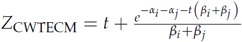.

With the change of variables *x*_*i*_ = *e*^−*α*_*i*_^, *y*_*i*_ = *e*^−*β*_*i*_^ the expected link probability and expected link weight have the same form found in Eq. 10, and are obtained by the first derivatives of the free energy with respect to the Lagrange multipliers *α*_*i*_ and *β*_*j*_ as follows:

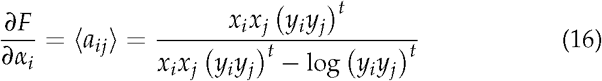

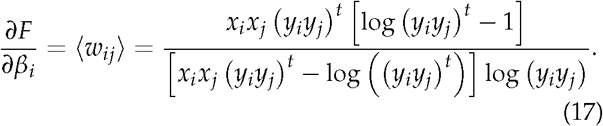

The expected degree and strengths are found by summing the link probability and the expected link weights over all remaining nodes:

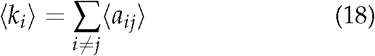

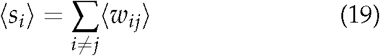

and at the optimal parameters 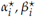, they equal their empirical counterparts 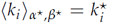 and 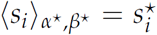. Similarly to the CWTECM, the optimal parameters can be found by maximization of a log-likelihood function that reads:

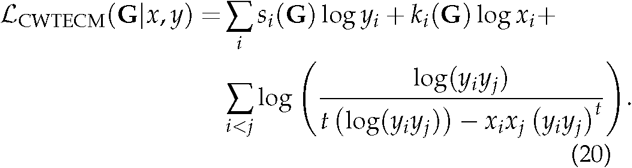

### 2.4. Spectral entropies

Classical maximum entropy methods are a tool for describing the ensemble of networks that show on average the same desired descriptor. However, when considering the problem of comparing two networks at all scales on a wide variety of metrics, classical maximum entropy methods fail in providing a statistically reliable tool, as one should design and solve a specific maximum entropy problem for every specific descriptor [16, 39]

However, it is possible to extend the maximum entropy approach to networks represented as quantum mechanical systems, by replacing the Shannon entropy with the von Neumann entropy [16, 19]:

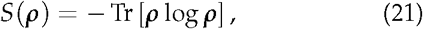

where ***ρ*** is the von Neumann density matrix, a Hermitian and positive definite matrix with unitary trace, that admits a spectral decomposition as:

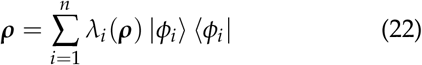

for an orthonormal basis {|*φ*_*i*_⟩} and eigenvalues *λ*_*i*_(***ρ***). By application of the maximum entropy principle where the networks in the ensemble are constrained to have on average the same Laplacian matrix, one finds that the role of classical probability in this case is replaced by the following density matrix:

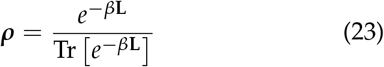

that describes the result of constraining the diffusion properties on the networks, and is in the form of a quantum Gibbs-Boltzmann distribution. The denominator *Z* = Tr [*e*^−*β***L**^] is the so-called partition function of the system (to be distinguished from *Z* in Eq. 4 of the null models), which can also be expressed as the sum of the eigenvalues of the matrix *e*^−*β***L**^ as follows:

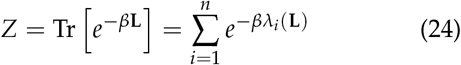

with *λ*_*i*_(**L**) the eigenvalues of the Laplacian **L**. The von Neumann density matrix defined in Eq. 23 is based on exponentially scaled eigenvalues of the graph Laplacian, and contains contributions at different scales of patterns in the network. Yet, the role of the Lagrangian multiplier *β* is far from trivial [19]. The *β* parameter can be interpreted as an inverse temperature (in classical statistical mechanics) or a normalized time [19] in modeling heat diffusion over the network. In the *β* → 0 limit, the density matrix can be expanded linearly as ***ρ*** ~ **I** − **L** and carries information about local connectivity patterns. On the other hand, for *β* → ∞, the diffusive behaviour is governed by the smallest non-zero eigenvalue of the Laplacian *λ*_2_, hence ***ρ*** ~ *e*^−*βλ*_2_^ |*φ*_2_⟩ ⟨*φ*_2_|, where |*φ*_2_⟩ is the eigenvector associated to *λ*_2_. This eigenvector is also called the Fiedler eigenvector and embodies the large scale structure of the graph [48].

The Laplacian spectrum encloses several critical topological properties of graphs [49–52]. For instance, the multiplicity of the zero eigenvalue corresponds to the number of connected components, while the multiplicity of each eigenvalue is related to graph symmetries [51–53], and the concept of expanders and isoperimetric number are connected to the first and second-largest eigenvalues [54, 55]. Moreover, the graph Laplacian appears often in the study of random walkers [56, 57], diffusion [58], combinatorics [59], and a large number of other applications [51, 59]. For this reason, the spectral entropy, which is ultimately based on Laplacian eigenvalues, describes a large number of typical properties of the network, aggregated in a single quantity.

For the sake of comparing two different networks represented by the density matrices ***ρ*** and ***σ*** here we use the notion of von Neumann relative entropy [16, 18] that reads:

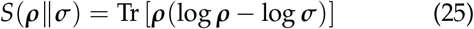

This definition encloses the concept of network similarity, as the relative entropy is a positive quantity, and is zero if and only if ***ρ*** = ***σ***. For this reason, and in the rest of this manuscript, we quantify the similarity between a network and its randomized counterpart by means of von Neumann relative entropy *S*(***ρ*** ‖ ***σ***). Additionally, it is straightforward [16, 19] to show that the minimum of relative entropy corresponds to the maximum of a log-likelihood functional, log 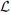, defined as

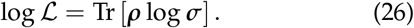

Similarly to Eq. 5, in the spectral framework it is possible to decompose the log-likelihood into the sum of Hamiltonian and free energy. This happens in equilibrium conditions where the density matrices ***ρ*** and ***σ*** are in the form of a Gibbs distribution like specified in Eq. 23. In particular, denoted ***ρ*** and ***σ*** as the density matrices of the empirical network **G\*** and of its randomized counterpart **G** with Laplacian **L**, respectively, the resulting spectral log-likelihood from Eq.26 takes the following form:

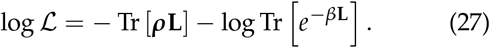

Here we compare networks with their randomized counterparts sampled from the maximum entropy models CWTERG and CWTECM using spectral entropies. We leverage spectral entropies, in order to resolve differences between networks at different scales governed by the diffusion parameter *β*. Indeed, graph models that preserve the local features (like nodal strength and degree) exhibit comparatively similar spectral entropies to those of the empirical network in the regime *β* → 0. Conversely, mesoscopic structures that cannot be modeled by solely constraining local features, will result in different von Neumann entropy *S* in the large *β* regime.

This concept is illustrated in Figure 2, where a highly regular network and its randomized counterpart are depicted, together with their von Neumann entropies, over a large range of the *β* parameter. The orange line in Figure 2A, describes the entropy *S* of the modular network. A large plateau indicates the tendency of a random walker to remain trapped in medium-size highly dense subset of nodes: its height is indicative of the logarithm of the number of modules *B*. On the other hand in the two extreme *β* regimes, entropy tends to its maximum or minimum attainable values, log *n* or log *C* respectively, where *C* is the number of weakly connected components. A degree-preserving randomized network has more uniform diffusion properties instead, as testified by the sharply falling blue dashed curve, where no specific mesoscopic pattern can be found. In the *β* → 0 limit, diffusion is only limited to the local neighborhood of nodes, hence, the orange and blue curves look similar. With a physical metaphor, random walkers spend more time at increasingly larger and isolated structures of the modular network, from small clusters to larger modules (Figure 2B). Finally, at steady-state *β* → ∞ both the two curves converge to the global structure, only specified by the number of connected components.

**Figure 2.**
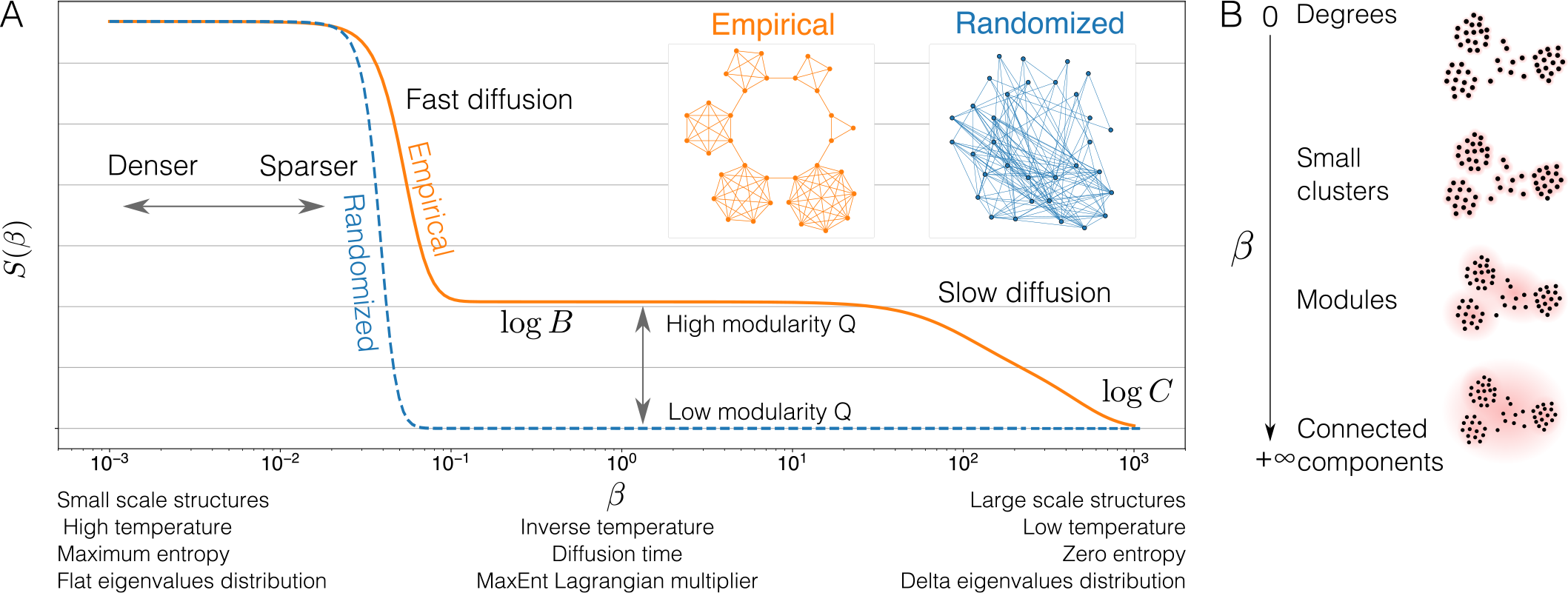
Von Neumann spectral entropy of a highly-ordered network (orange), and its randomized, degree preserving counterpart (blue). Panel A. For small values of *β* the spectral entropy reaches its maximum value *S* = log *n*, while in the large *β* limit it tends to the logarithm of the number of connected components (log *c*, zero for weakly connected graphs with *c* = 1 components). Intermediate values of *β* highlight mesoscopic structures. The height of the plateau is related to the overall modularity of the network, while its positioning on the horizontal axis depends on network links density. Differently from the highly regular ring of cliques (orange), the randomized network (blue) shows no structure at all scales, hence its von Neumann entropy decreases rapidly. Panel B shows that low *β* correspond to local features while large *β* describes large scale features.

The concepts and models exposed in this section are implemented in the open source package *networkqit*, written in Python.

## 3. DATA AND PREPROCESSING

For the purpose of this study we selected publicly available resting-state empirical networks as benchmarks to test this new theoretical framework.

### 3.1. Resting-state network

To evaluate the effects of different thresholding on the network structure, we have chosen a resting-state network computed as a group average of 27 healthy volunteers (mean age 24 yrs.) and described in Ref. [60], alongside with the ethical statements. This functional connectivity network is a popular benchmark for testing graph-theoretical methods, and was chosen to enable direct comparison with previous literature. The connectivity matrix is available at Ref [61]. Functional data were acquired with a Siemens Tim Trio 3T scanner, with a TR=2 s, TE=31 ms, voxel size 3.5 × 3.5 × 3 mm, for a total of 153 volumes recorded for 5 minutes and 6 seconds. Regional time-series were extracted for 638 nodes using Crossley’s parcellation scheme [60], head rotations and translations together with their derivatives and mean cerebrospinal fluid time series were regressed and band-passed (0.01–0.1 Hz). The functional connectivity matrix was derived computing pairwise Pearson correlations, normalized by the Fisher transform, and finally across subjects. The network corresponds to the unthresholded version made publicly available through the Brain Connectivity Toolbox (BCT, [62]). To assess the effect of thresholding, we applied a range of different absolute thresholds, from *t* = 0.1, to the point where the network remains fully connected. Above this threshold, nodes start detaching from the main largest connected component, reflecting the hierarchy of modules comprised in the network [63]. Here, absolute thresholding corresponds to the removal of all edges with weight *w*_*ij*_ < *t* where *t* is a real positive number. The point where the network breaks apart is dubbed percolation point. Thus, with the term percolation threshold we mean the highest value of absolute threshold *t*\* such that the undirected network remains connected, i.e. it comprises a single connected component. The percolation analysis of the Crossley network is shown in Figure 3.

**Figure 3.**
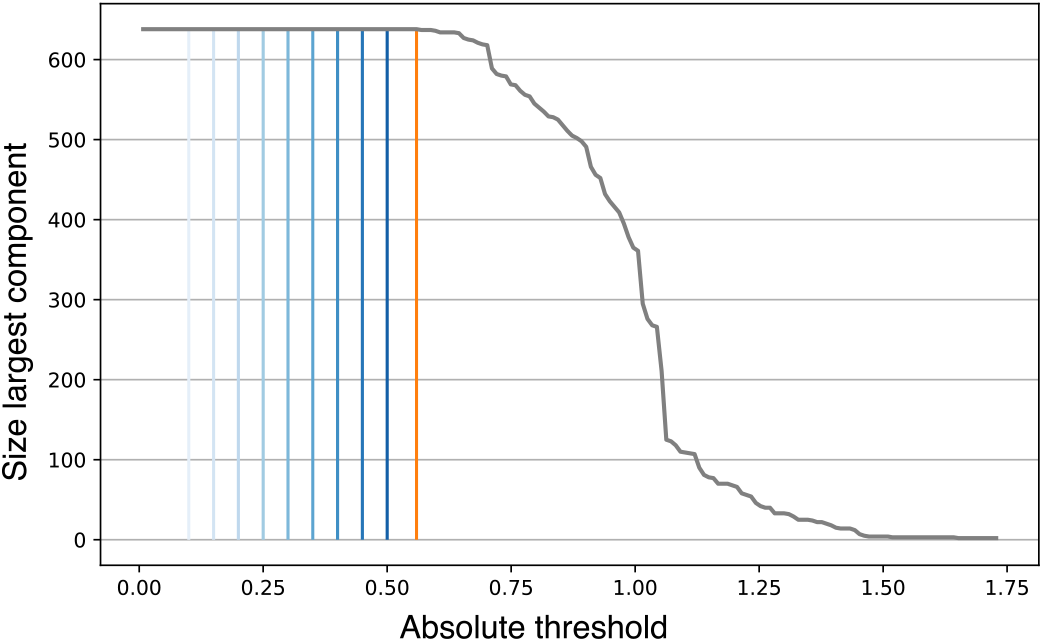
Percolation analysis for the healthy resting state dataset of 638 nodes, described in Crossley et.al [60]. The gray curve indicates the size of the largest connected component as a function of the threshold. The blue lines correspond to threshold values from 0.1 to 0.5, the orange line is the percolation threshold, where the largest connected component starts breaking apart.

### 3.2. Motion and motion correction

To study the effects of motion, we selected neuroimaging data from the MPI - Leipzig Study for Mind-Body-Emotion Interactions project (LEMON, [64]), obtained from the OpenfMRI database, accession number *ds000221*. Ethical statements are present in the original references by the groups who performed the experiments. Given the growing concern in the neuroimaging literature regarding the effects of head motion on resting-state functional connectivity data [65], and the impact of different motion-correction techniques [31], we decided to explore the performance of the two null models over matrices containing different degrees of motion and different preprocessing pipelines. Indeed, according to recent findings, even very small head movements (0.2 mm) can substantially affect functional connectivity networks, increasing spurious correlations and altering its underlying topology [29, 65]. The use of this large data-base has enabled us to select subgroups of subjects with different levels of motion and adequate size for a significant between group comparison.

From this dataset, participants were selected according to the age range; only participants ranging from 20 to 30 y.o. were included in our study, to avoid age effects in subsequent analyses. All MRI data were acquired with a 3T scanner (Magnetom Verio, Siemens Healthcare, Erlangen, Germany), with the following parameters TR = 1.4 s, TE = 39.4 ms, for a total of 657 volumes, resulting in 15 minutes and 30 seconds of recording. A total of 117 subjects were selected. Structural and functional images were preprocessed with FSL (v 5.0) [66]. High-resolution structural images were registered to the MNI template and segmented (fast segmentation), separating white matter and ventricles masks. Functional preprocessing included motion correction and realignment (mcflirt), coregistration to the structural image using boundary based registration (BBR) and then normalized to the MNI template. For each participant, we extracted regional mean time series from 638 parceled areas, based on the same template employed for the other networks already addressed in the present study. A Butterworth bandpass filter between 0.01 and 0.1 Hz was applied to all the time series.

For the purposes of the study, all participants were divided into three groups according to their degree of motion (Low, Medium, and high-motion), measured as the proportion of outlier volumes present within the time series. To evaluate the motion level of each subject, Framewise Displacement (FD) was computed according to Power [65]. Timepoints were flagged as outliers affected by motion when FD > 0.3mm. Criteria for group subdivision, decided after careful inspection of the data, were the following:

- Low-motion (*N* = 39): less than 1% data affected.
- Medium-motion (*N* = 39): data affected between 1% and 5%.
- High-motion (*N* = 39): more than 5% data affected.

The three groups were balanced for age and sex, but different for in-scanner motion.

Based on the growing debate related to the best noise-correction technique to apply on resting-state data, we tested two different pipelines, plus one pipeline where no de-noising strategy was applied. We selected and analyzed the results on the following pipelines:

- P0: no motion-correction technique applied beside rigid image realignment carried out with *mcflirt* [67];
- FIX: based on the FMRIB trained classifier of Independent Component Analysis, components related to noise (FIX, [68]), extracted from single-subjects time series;
- 9P: a common method that requires the regression of different factors, such as 6 movement parameters, the average signal extracted from white matter and cerebrospinal fluid, plus the regression of the global signal (GSR), measured as the average of all the voxels of the brain extracted from subject-specific brain masks.

Before the regression of all the confound parameters from subjects’ time series, a Butterworth bandpass filter of 0.01 and 0.1 Hz was applied to all the regressors, avoiding reintroduction of signal related to nuisance co-variates [69].

Altogether, we specifically selected pipelines based on different principles. One strategy relies on independent components classification (FIX), the second includes the regression of the global signal (9P), a controversial practice. As a reference, for the simple evaluation of pure effects of motion over the architecture of the functional network, we considered a pipeline where only the mandatory image preprocessing steps (realignment, normalization, coregistration, filtering) have been applied (P0).

Differences among groups in terms of connectivity strength were measured by means of simple t-tests. Overall functional connectivity strength in every network was addressed as the mean of all positive links [10]. From an effective pipeline we would expect a reduction in the differences induced by motion in the three groups. At the same time, we would expect that the attenuation of these differences would not alter the topological structure of the functional network. Conversely, an excessively aggressive motion-correction approach may also remove genuine correlations, thus erasing large-scale network structure.

## 4. RESULTS AND DISCUSSION

### 4.1. Null model fitting

As a first application of the models and framework exposed in the previous sections, in Figure 4 we report the results of the maximum likelihood estimation of the Continuous Weighted Enhanced Configuration and of the CWTERG models in the network of healthy subjects (Crossley, [60]) described in section 3 3.1.

**Figure 4.**
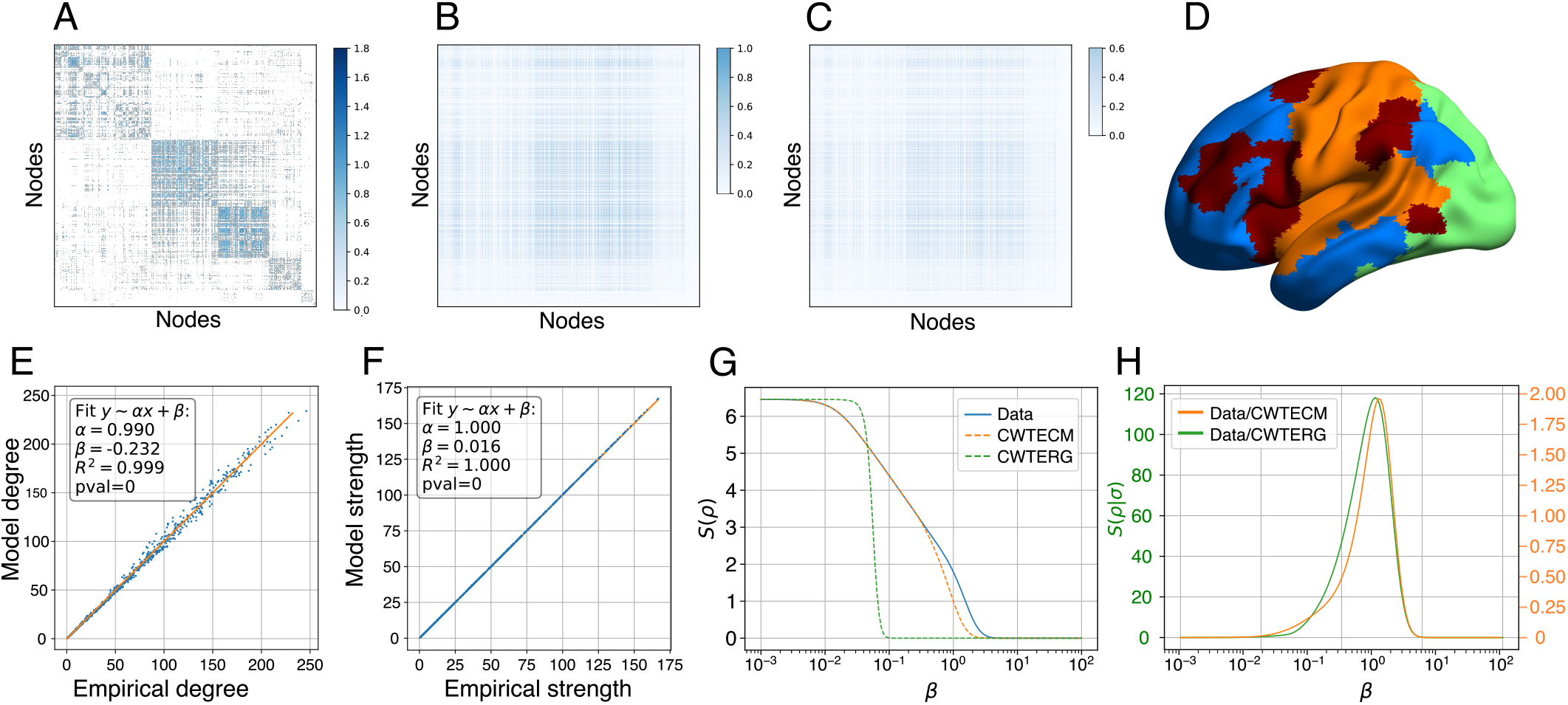
Continuous Weighted Thresholded Enhanced Configuration Model fitted on a real functional network. Panel A. The empirical functional connectivity matrix, thresholded at percolation with rows and columns reorganized by the maximum modularity community structure, to highlight the community structure as overlaid on a brain in Panel D. Panels B,C. the link probability and expected link weights as from Eqs. 16,17. Panels E,F. the reconstructed degrees and strength as sums over the rows of the link probabilities and expected weights matrices. On the horizontal axis the empirical degrees (strengths), on the vertical axis the model degrees (strengths): *R*^2^ and regression slopes are shown as inset. Panel G. the spectral entropy of the network (blue line), CWTECM null model (orange dashed line) and CWTERG null model. Panel H. relative entropy of the network with respect to CWTECM (orange line) and CWTERG (green line). Link probability and expected weights for the CWTERG are not shown, as they are scalar numbers.

The fully-dense network was thresholded at percolation level, corresponding to a links cut-off value of *t* = 0.55 (Figure 4A). The maximization of log-likelihood resulted in the optimal parameters *x*\*, *y*\* defining the link probability *p*_*ij*_ (Fig. 4B) and the expected link weight ⟨*w*_*ij*_⟩ (Figure 4C), as defined in equations 16 and 17.

The empirical degree and strength sequences are depicted in Fig. 4E,F compared to the reconstructed degrees and strengths from the model. In order to quantitatively assess the level of reliability of the estimated parameters, we fitted a linear model between empirical and model networks. The results are shown in the insets of Panels E and F of Figure 4. The regression slope of the degrees is very close to identity *α* = 0.99 with a very high *R*^2^ coefficient. The strength reconstruction is also very accurate with a regression slope of 1. Panels G and H of Figure 4 demonstrate the difference between the optimal models (CTWERG and CWTECM) in terms of spectral entropies curves and relative entropies as a function of the scale parameter *β*. The spectral entropy of the CTWERG model fits that of the empirical network only at local scale, and drops rapidly for larger betas. Conversely, the spectral entropy of the optimal CTWECM closely matches its empirical counterpart for a wide range of beta, with the exception of the largest scales. This behaviour indicates that fixing the degree and strength sequence represents a strong constraint, which determines the network structure at the meso-scale. Spectral entropy at large scale reflects the network’s modular organization, which cannot be accounted for by local constraints.

### 4.2. Effects of thresholding

Here we use these null models within the theoretical framework of spectral entropies to explore the effects of network sparsification on the structure of functional connectivity networks.

Specifically, we applied different levels of absolute thresholds to the empirical network and to the models (from *t* = 0.10 to the point where the network breaks apart). Hence, we computed the spectral entropies of thresholded networks and corresponding null models, for different values of *β*. We then used relative entropy to quantify the information-theoretic distance as a function of threshold. The hypothesis is that the distance between the empirical network and its maximally random counterpart depends on the sparsification threshold, and that there may exist an optimal threshold value that maximizes this distance, striking the optimal balance between removal of spurious correlations and undesirable suppression of structural information.

The results are summarized in Figure 5, which shows spectral and relative entropies for the empirical network and both null models at various threshold levels. Firstly, we observe that at lower threshold levels (depicted in light blue, Fig. 5) the relative entropy decreases sharply as a function of *β*, a result of faster diffusion time. Indeed, lower threshold levels correspond to denser networks, and consequently faster diffusion. Decreasing network density results in a right-shift of spectral entropy curves for all networks, as shown in Figure 5A,C. However, the effects of thresholding are different in the empirical network and in its null models, a result of the structure of the functional connectivity network that is not accounted for by its randomized counterparts.

**Figure 5.**
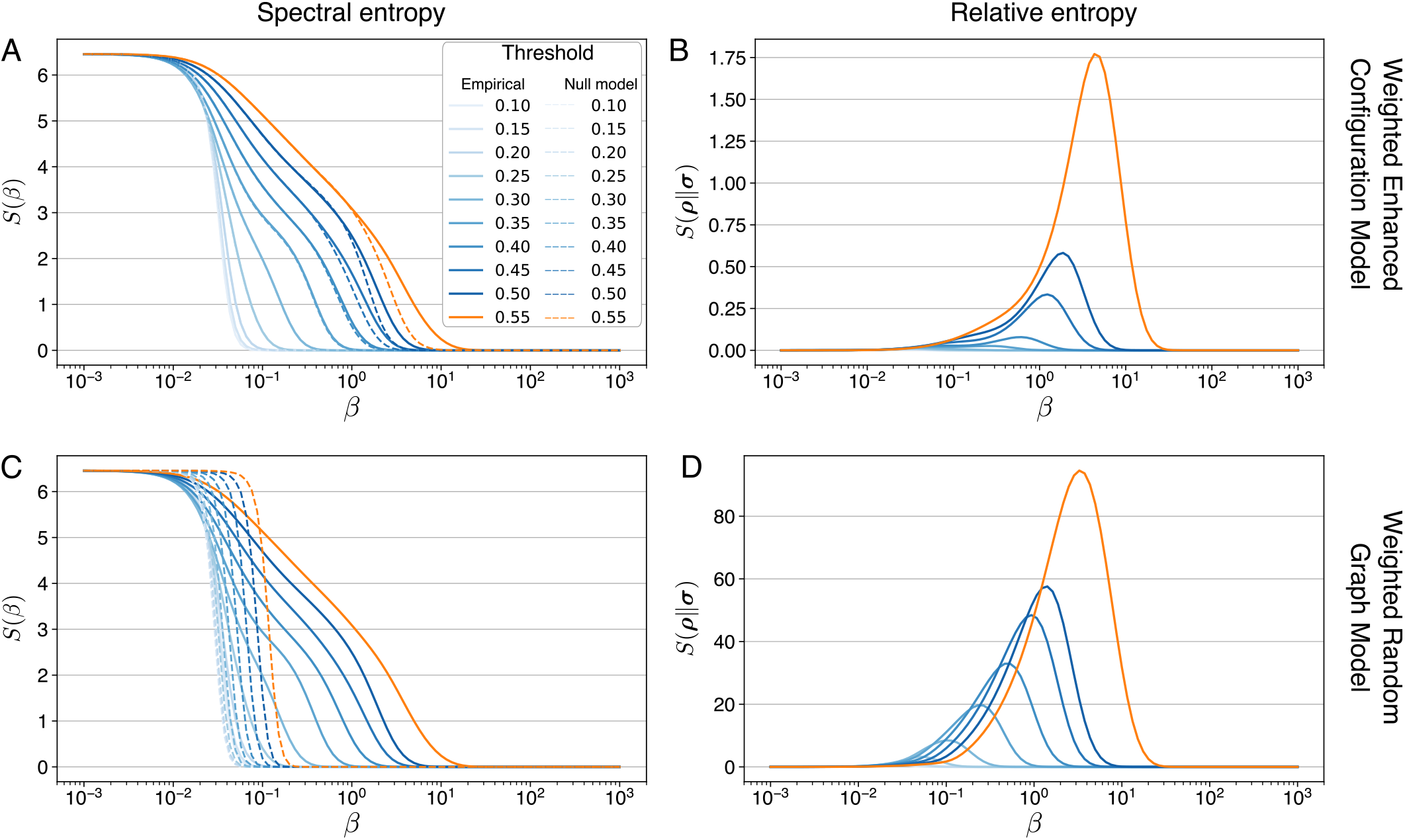
Spectral entropies and relative entropy of the Crossley’s functional connectivity network compared to its randomized counterparts. Blue shaded lines represent networks thresholded at absolute values from 0.1 to 0.5. Orange lines denote the network at percolation. Solid lines are referred to the empirical network. Dashed lines are referred to the randomized models. Panels A,B show the results with respect to the CWTECM model. Panels C,D show the results with respect to the CWTERG model.

Indeed, large-scale structures of the empirical network emerge at higher thresholds (darker blue), as reflected in Figure 5A by the presence of “information shoulders” i.e., plateaus in the spectral entropy curve. This phenomenon is not equally present in the two null models. Should the thresholding procedure highlight mesoscopic structures only accounted by local constraints, we would expect similar high values of *S* on both the thresholded random counterparts of the empirical graph. However, the CWTERG shows no indications of a high-level organization at any threshold, as seen by the sharply falling entropy within a very small range of *β* (Figure 5C). Indeed, as previously demonstrated, the CWTERG destroys network structure by completely shuffling nodes’ neighborhoods. As a result, the diffusion process rapidly spans the whole network, as every node has uniform probability of being connected to every other node.

On the other hand, the spectral entropy of the CWTECM closely corresponds to the one of the empirical network over a broad range of *β* values. Significant differences only appear at large scales for increasing thresholds. This result shows that degree and strength sequence constrain local and medium-scale structures. In accordance with the results of reference [23], we observed that the large-scale community structure is the only feature that is not accounted for by local properties.

This same phenomenon is also reported in Figure 5B,D where the relative entropies are shown for both models. The relative entropy for the CWTERG attains a higher maximum at slightly smaller values of *β* than for the CWTECM. Intuitively, it takes less time for a random walker to explore a random network than a complex network where modules and local structures may hamper the diffusion process. Moreover, for both cases, relative entropies accentuate the effects of thresholding, as they increase with increasing sparsity level, peaking around the percolation point, just before the network starts breaking apart.

The results in Figure 5 demonstrate that, in both cases, the maximal spectral entropy difference of the empirical network from its null model is found around the percolation threshold.

Taken together, these results demonstrate the beneficial effects of sparsification to enhance and retrieve the network’s modular structure. Importantly, we find that this effect is maximal at percolation threshold, above which the network starts breaking apart into unconnected sub-units. At low threshold levels, the empirical network is remarkably close to a random network with similar local features. Only at higher thresholds its meso- and large-scale structure emerges, and difference from the null models become apparent. Interestingly, the presence of an optimal threshold appears to be independent of the specific null model. These results are consistent with previous empirical findings in model networks [70] and provide a theoretical foundation to the use of sparsification methods to study the large-scale topology of functional connectivity networks.

### 4.3. Effects of motion and motion correction

We have also applied the concept of an information-based comparison with null models to investigate the effects of motion and motion correction pipelines typically used in fMRI studies. A major debate in the resting-state functional connectivity community concerns the effects that different preprocessing pipelines can have over the functional time series [31, 65]. Here, our goal is to investigate to what extent motion could render the network more or less similar to its random counterpart, and the efficacy of different pipelines and thresholding procedures in mitigating the effects of motion. We used the same atlas with 638 parcels that was used in the previous examples, and we applied the same model fitting and comparison techniques to resting-state dataset with different degrees of motion and different motion-correction techniques (see Data and Preprocessing, section 3). Specifically, we considered three pipelines: P0 with no motion correction, a second pipeline based on FIX [68], and 9P, a pipeline that includes global signal regression [31]. We applied these pipelines on three motion groups: low, medium, and high-motion (see Data and Preprocessing, section 3). These three groups are defined on the basis of Framewise Displacement (FD), a metric commonly used to evaluate the amount of head motion in rsFC [28], that is computed as the sum of the absolute values of the derivatives of the six motion parameters. Power and colleagues [71] showed that even small head movements (FD > 0.15 mm) could cause significant changes affecting all voxels in the brain. These movements can be visually identified utilizing so called *grayplots*, as shown in Figure 6. Grayplots depict the signal intensity of every voxel in the brain over time. Here, we report three examples of how the magnitude of head movements impacts the whole time series. In Figure 6 one can appreciate the effects of motion as abrupt changes of voxel intensities in correspondence of head movements. These artifacts give rise to spurious correlations at different scales.

**Figure 6.**
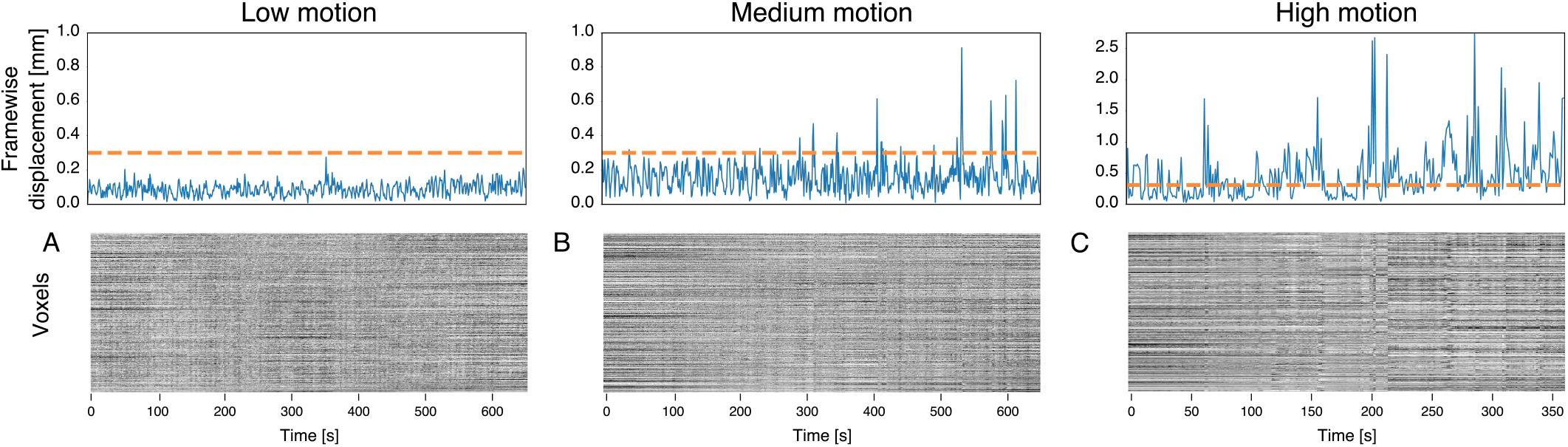
Framewise displacements of the three motion groups together with the gray plots of the fMRI scans. The orange line at FD 0.3mm represents the limit of outliers.

The three groups here evaluated are perfectly balanced for age and gender, with exactly the same acquisition procedures, but different for in-scanner motion. Thus, at the group level, we would expect these participants to share same global functional connectivity characteristics. Possible differences should be driven by the presence of motion.

Firstly, we evaluated the effects of motion on functional connectivity strength. In line with previous reports [31], we observed a substantial increase in functional connectivity induced by motion in P0 (Figure 7). Figure 7A shows that the distribution of link weights for the medium and high-motion groups is right-shifted compared to the low groups, reflecting higher correlation strength. At the subject level this shift is highly significant across all groups (medium > low: *p* < 10^−5^; high > low: *p* < 10^−4^). Through the application of specific denoising strategies we sought to investigate to what degree this spurious difference in functional strength between groups can be reduced. The histograms despicted in panels B and C of Figure 7 show that both pipelines appear to significantly decrease the differences in edge-weight distribution at the group level across different motion conditions. Specifically, the pipeline based on independent components classification (FIX) substantially reduces the right shift of the medium and high-motion groups that was apparent with P0. Importantly, in this condition the edge-weight distribution of the medium-motion group almost completely overlaps with the low-motion curve. Yet, at the individual level the functional connectivity strength, measured as the mean of all positive edges in the graph, shows statistical difference (medium > low: *p* = 0.007). In contrast, the histogram representing the edge-weight distribution extracted from the high-motion group still presents a highly significant right shift reflecting higher functional strength (high>low: *p* < 10^−3^). A different pattern is revealed in the strength distribution after the application of GSR. In this case, all curves are highly overlapping, indicating similar functional connectivity across the three groups. This is revealed by the lack of significant differences at the individual level in edge strength (high > low: *p* = 0.8; medium > low: *p* = 0.48). As a matter of concern, all three distributions are centered around zero. Indeed, after GSR the number of negative correlations dramatically increases, involving almost half of the edges within the network. These observations are in line with previous reports and concerns related to the controversial application of this denoising approach [31].

**Figure 7.**
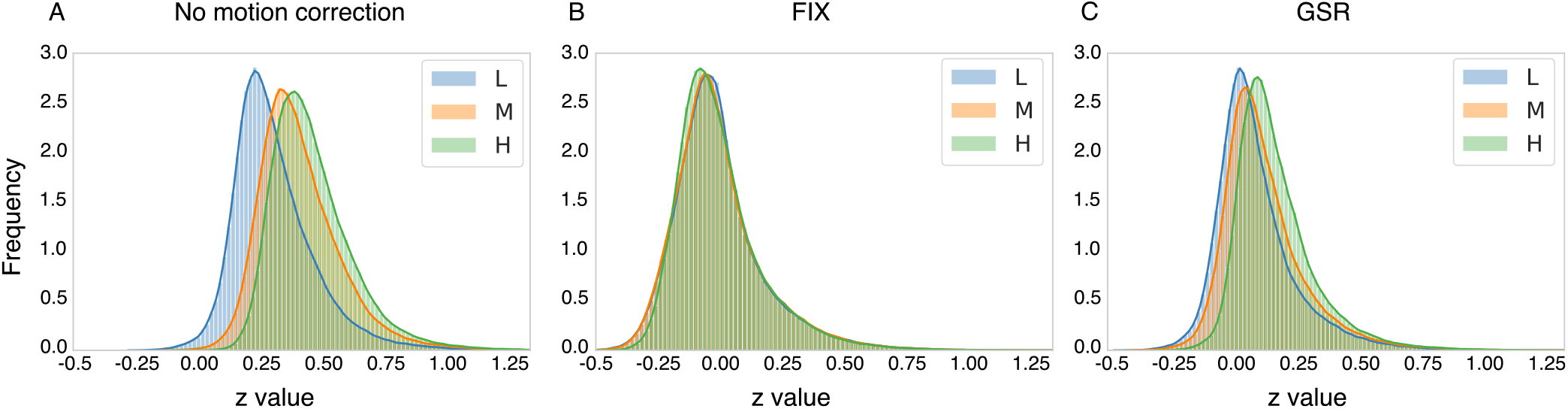
Effects of motion and commonly applied motion-correction techniques over the distribution of functional connectivity strength. Panel A depicts effects of motion as assessed by means of a pipeline where no motion correction strategies has been applied (P0). As a consequence of motion, we observe strong changes in the functional connectivity strength across the three groups (medium>low: *p* < 10^−4^; high¿low: *p* < 10^−5^). Panel B represents the effects of the application of FIX over edge-weight distribution. Differences among groups are still present but attenuated (medium*>*low: *p* = 0.007; high>low=*p* < 10^−3^). Panel C shows the effects of application of GSR. Differences among groups are not present (medium>low: *p* = 0.48; high>low: p=0.8).

In Figure 8 we show the spectral entropy curves and relative entropies for the three pipelines considered. In light of the previous findings, we present here only the results related to the CWTERG, as the constraints imposed by the CWTECM also affect larger scales, and may reduce sensitivity to the effects of motion at a mesoscopic level. Panels A,B and C show the von Neumann entropy curves of the differently pre-processed resting-state networks across three degrees of motion. In this specific case, we applied a single threshold, namely the lowest absolute threshold that guarantees connectedness in all three motion groups within the same pipeline (*t* = 0.44 for P0, *t* = 0.29 for GSR, *t* = 0.25 for FIX). We can observe that, when considered at the same absolute threshold, the low-motion group always shows higher spectral entropy across the entire *β* range. This is especially evident in the P0 pipeline. It appears, indeed, that movement artifacts significantly affect the mesoscopic patterns within the empirical network. This trend is confirmed by the smaller entropy values of the high-motion group compared to the medium and low-motion groups, which is observed across all analysis pipelines. In Figure 8A, this point is further highlighted by the lack of a clear-cut modular structure in both the medium and high-motion groups, whereas a shoulder present at medium scales for the low-motion group denotes a different degree of inter-modular density. A similar trend suggests that head movements tend to make the network closer to its null model, i.e. more random. Popular correction techniques mitigate this confounding effect, decoupling functional connectivity and motion.

**Figure 8.**
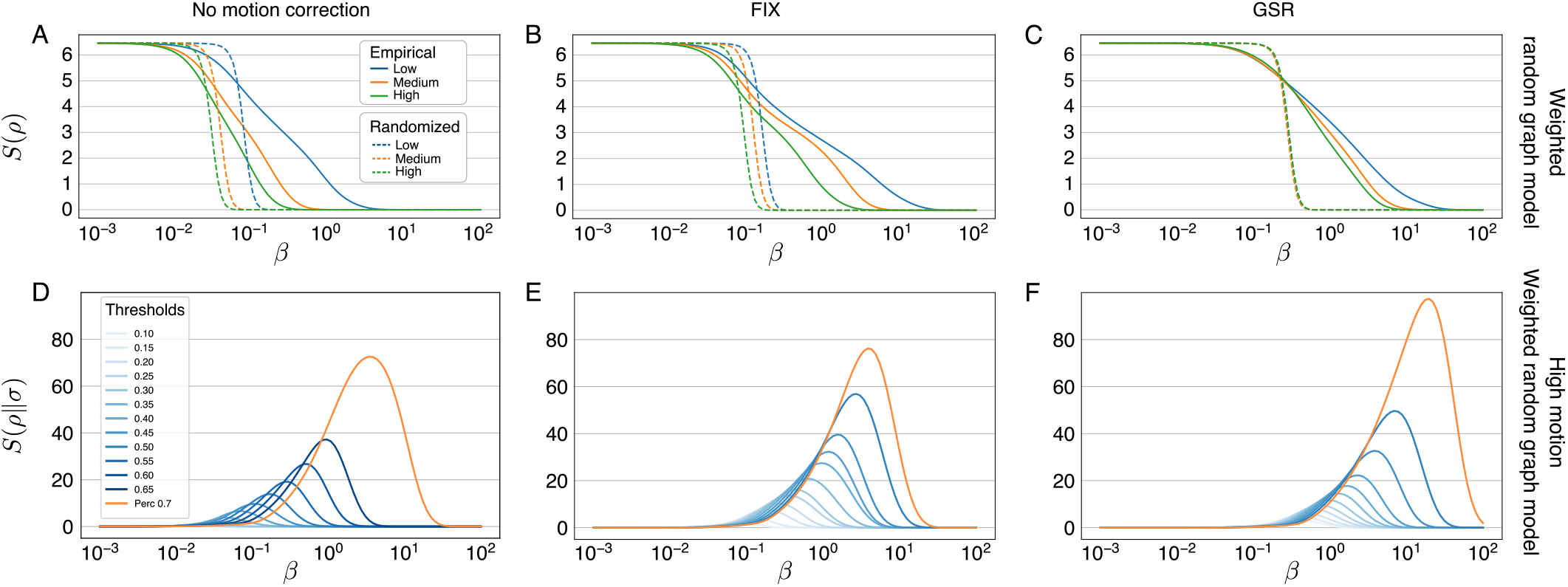
Panels A,B,C show the spectral entropies of networks for the pipelines 0,FIX and GSR (solid lines), together with their randomized counterpart (CWTERG, dashed lines) over all motion groups. The relative entropies of networks from the high-motion group are shown in panels D, E, F where the blue shades correspond to increasing absolute thresholds, while the orange lines correspond to percolation threshold, which has the maximum relative entropy at large scales.

Panels B,C of Figure 8 show spectral entropy curves for the pipelines FIX and GSR, respectively. As already discussed, both pipelines importantly reduce the difference in spectral entropy between the three groups. It is noteworthy that the application of the FIX pipeline in Figure 8B highlights the manifest presence of more prominent shoulders in the entropy curve in all groups, again a signature of mesoscopic organization. From this view, the cleanup of resting-state data through independent component analysis appears to emphasize the global structure of the network, in presence of head movements. The same cannot be appreciated in the groups preprocessed with a GSR pipeline: differences in spectral entropies are reduced, but no clear large-scale structure seems to emerge from these curves.

From the relative entropies generated over several different absolute thresholds, we can appreciate a strong effect related to the sparsification procedure. In Panels D, E, F of Figure 8, the relative entropies of the high-motion groups for all pipelines and their respective null models are presented. In the interest of space, we report here only the high-motion group, which is more affected by head movements and shows more evidently the beneficial effects of the application of different preprocessing pipelines and thresholds. With the application of increasing thresholds, the distance of the empirical network from its random counterpart with same density gets larger, and it reaches maximum at percolation, despite the presence of motion and independently of the pipeline applied. Specifically, we can observe a higher relative entropy at percolation for the pipeline based on GSR (Figure 8F). This preprocessing technique notably benefits from the thresholding procedure, considering the substantial difference between the maximum relative entropy attained at percolation and its values for denser networks. In line with previous studies [31], the main effect of GSR is an increase in network modularity, mirrored by greater values of relative entropies at large scales, suggesting a well organized high-order architecture. Yet, the lack of an “information shoulder” in the spectral entropy curve suggests the presence of a more uniform structure, with similar intra-modular density across different communities, and similar size of the modules. Importantly, we note that thresholding emphasizes meso- and large-scale structure in combination with FIX (Fig. 8F), but also in the absence of any motion correction (Fig. 8D). Indeed, network sparsification appears to have a large effect per se, even for P0, in separating the empirical network form its null model. Interestingly, relative entropy between empirical network and null model increases with the threshold, and reaches its maximum at percolation. This last observation further supports the application of a thresholding procedure, in contrast with recent literature that suggests that sparsification should be avoided [24–27]. Our results demonstrate the importance of application of a threshold, irrespectively of the pre-processing pipeline.

## 5. CONCLUSION

The nature of resting state functional MRI networks based on pairwise association measures, like the Pearson correlation, is of a dense square matrix. Several experimental factors are involved in shaping the properties of these matrices and no consensus exists in the literature on the best practice for the definition and processing of these matrices and the associated connectivity graphs. In the present work we have introduced a novel theoretical framework than contributes to shed light on several open issues in the analysis of brain functional connectivity networks. Firstly, we define the CWTERG and CWTECM null models, which enable extension of the maximum entropy random graph formalism to networks with threshold and real positive weights, as those encountered in fMRI. Secondly, we have shown that the spectral entropies framework can be applied to the differences of networks with respect to their random versions from local to global scales.

Leveraging this new approach, we studied the effects of thresholding procedures and motion-correction pipelines. The application of a threshold to resting-state networks is a contentious step debated in the field.

Here, by means of advanced information theory tools, we found that complete functional connectivity networks present a high degree of randomness, due to the contribution of spurious correlations to weak links, that conceals their large scale structure. Sparsification of the network is an essential step to differentiate networks from their null model and is highly beneficial to study the large scale architecture of real-world networks.

Further, we demonstrated from first principles the existence of an optimal thresholding point, where the empirical network is maximally distant from random. Specifically, we found that application of a percolation threshold strikes the optimal balance between the removal of spurious connections and genuine information, thus maximizing the information that can be extracted from the system. The importance of sparsification can also be appreciated through the evaluation of the effects of motion and different preprocessing pipelines. Motion increases randomness and reduces spectral entropies across the whole *β* domain, bringing the network closer to its random counterpart. The effects of motion are mitigated by popular motion-correction approaches. However, we found that network sparsification has a beneficial effect irrespectively of the specific denoising strategy applied, and that the percolation threshold maximizes the distance of the empirical network from its randomized counterpart.

In conclusion, the novel framework of spectral maximum entropy networks provides a new and powerful approach that significantly extends the repertoire of tools for the study of functional connectivity networks at multiple scales.

## 5.1. Author contributions

C.N. developed the mathematical methods and software. C.N., G.F analyzed the data. G.F. collected and processed the neuroimaging data. C.N., G.F., L.M., A.B. wrote the paper. All authors reviewed the manuscript.

## 5.3. Acknowledgments

The authors wish to thank Dr. Tiziano Squartini and Prof. Diego Garlaschelli for useful methodological discussions. The authors also thank Prof. Edward Bullmore and Prof. Nicholas Crossley for providing data and templates. This project has received funding from the European Union’s Horizon 2020 Research and Innovation Program under grant agreement 668863, SyBil-AA. L. Minati gratefully acknowledges employment and research funding by the World Research Hub Initiative (WRHI), Institute of Innovative Research (IIR), Tokyo Institute of Technology, Tokyo, Japan.

